# Cholera in Internally Displaced Persons Camps in Borno State—Nigeria, 2017: A qualitative study of the multi-sectorial emergency response to stop the spread of the outbreak

**DOI:** 10.1101/599787

**Authors:** Moise C Ngwa, Alemu Wondimagegnehu, Ifeanyi Okudo, Collins Owili, Uzoma Ugochukwu, Clement Peter, Isabelle Devaux, Lorenzo Pezzoli, Chikwe Ihekweazu, David A Sack

## Abstract

**Introduction/Background:** In August 2017, a cholera outbreak started in Muna Garage IDPs camp, Borno State-Nigeria, and >5000 cases occurred in six local government areas. This qualitative study evaluated perspectives about the emergency response to this outbreak.

**Methods/Findings:** We conducted 39 key informant interviews and focused group discussions, and reviewed 21 documents with participants involved with surveillance, water-sanitation-hygiene, case management, oral cholera vaccine, communications, logistics, and coordination. Qualitative data analysis used thematic techniques comprising key-words-in-context, word-repetition, and key-sector-terms.

Authorities were alerted quickly, but outbreak declaration took 12 days due to a 10 day delay waiting for culture confirmation. Outbreak investigation revealed several potential transmission channels, but a leaking latrine around the index cases’ house was not repaired for >7 days.

Use of chlorine disinfectant was initially not accepted by the community due to rumors that it would sterilize women. This could have been avoided with improved community consultation. Initially, key messages were communicated in Hausa, although ‘Kanuri’ was the primary language; later this was corrected. Planning would have benefited using exercise drills to identify weaknesses, and inventory sharing to avoid stock outs.

The response by the Rural Water Supply and Sanitation Agency was perceived to be slow and an increased risk from Eid El Jabir festival with increased movement and food sharing was not recognized. Case management was provided at treatment centers, but some partners were concerned that their work was recognized asking, “who gets the glory and the data?”

OCV was provided to nearly one million people and it distribution benefited from a robust polio vaccine structure; however, logistical problems related to payment of staff needed resolution.

Initial coordination was thought to be slow, but improved by activating an Emergency Operations Centre. The Borno Ministry of Health used an Incident Management System to coordinate multi-sectoral response activities.

These were informed by daily reviews of epi curves and geo-coordinate maps. The synergy between partners and government improved when each recognized the government’s leadership role.

**Conclusions/Significance:** Despite a timely alert of the outbreak, the delayed declaration led to a slowed initial response, but this improved during the course of the outbreak. OCV distribution was efficient and benefited from the OPV infrastructure. Improvements in laboratory capacity are urgently needed.

**Author Summary:** In August 2017, a cholera outbreak started in the Muna Garage Internally Displaced Persons (IPDs) camp in Borno State, Nigeria. By October, it appeared in six local government areas with a total of 5,340 cases reported including 61 deaths. We evaluated the perspectives of the emergency response by the government of Nigeria and implementing partners to stop the outbreak. We conducted 39 interviews and group discussions and also studied 21 documents related to the outbreak response. We found that epidemiologic surveillance timely alerted the health authorities about the outbreak, but the outbreak was declared 12 days later, awaiting for culture confirmation. This led to delays in the initial response. We also observed that conditions in the IDPs camps like overflowing latrines, overcrowding, and open defecation were highly favorable to cholera transmission. Improved IDP camp conditions are needed to prevent cholera and other water born infections and strengthened laboratory capacity is needed to enable a more rapid response.

## Introduction

Cholera is an infection of the intestines caused by the bacterium *Vibrio cholerae*. The infection is usually transmitted by consumption of contaminated food or water. It can lead to severe watery diarrhea, dehydration and in some cases, death within a few hours. The main stays of cholera prevention include provision of potable water, adequate sanitation, and proper hygiene. Despite these well-known prevention strategies, cholera claims an estimated 1.3 to 4.0 million cases annually, and 21,000 to 143,000 deaths worldwide [1].

Cholera was first reported in Nigeria (Fig 1A) in 1970 [2]. Between 1991 and 2018 a total of 321,148 cases and 18,644 deaths with 5.8% case fatality ratio (CFR) were reported across Nigeria [2, 3]. In 2010, Local Government Areas (LGAs) in Borno State (Fig 1A) reported an outbreak that grew in magnitude and spread to include 21,111 cases (CFR 5.1%) [4]. In August 2017, another cholera outbreak started in Borno State in the Muna Garage camp for Internally Displaced Persons (IDP) camp in Jere LGA (Fig 1B). By September 2017 (Fig 2), it spread to five other LGAs (Fig 1B). A total of 5,340 cases were reported with 61 deaths (CFR 1.14%). In response, the Nigeria Centre for Disease Control (NCDC), National Primary Health Care Development Agency (NPHCDA), Borno Ministry of Health (MOH) together with implementing partners carried out comprehensive cholera control measures including Oral Cholera Vaccine (OCV) to combat the outbreak. As part of Monitoring and Evaluation to inform future emergency response effort, the question arises what are the perspectives of government and partners about the outbreak response? This study addresses this question using a cross-sectional qualitative research design to evaluate the experience of the various agencies involved in the outbreak response. The study was designed to inform decision making in organizing emergency responses to future cholera outbreaks in Nigeria.

**Fig 1.**
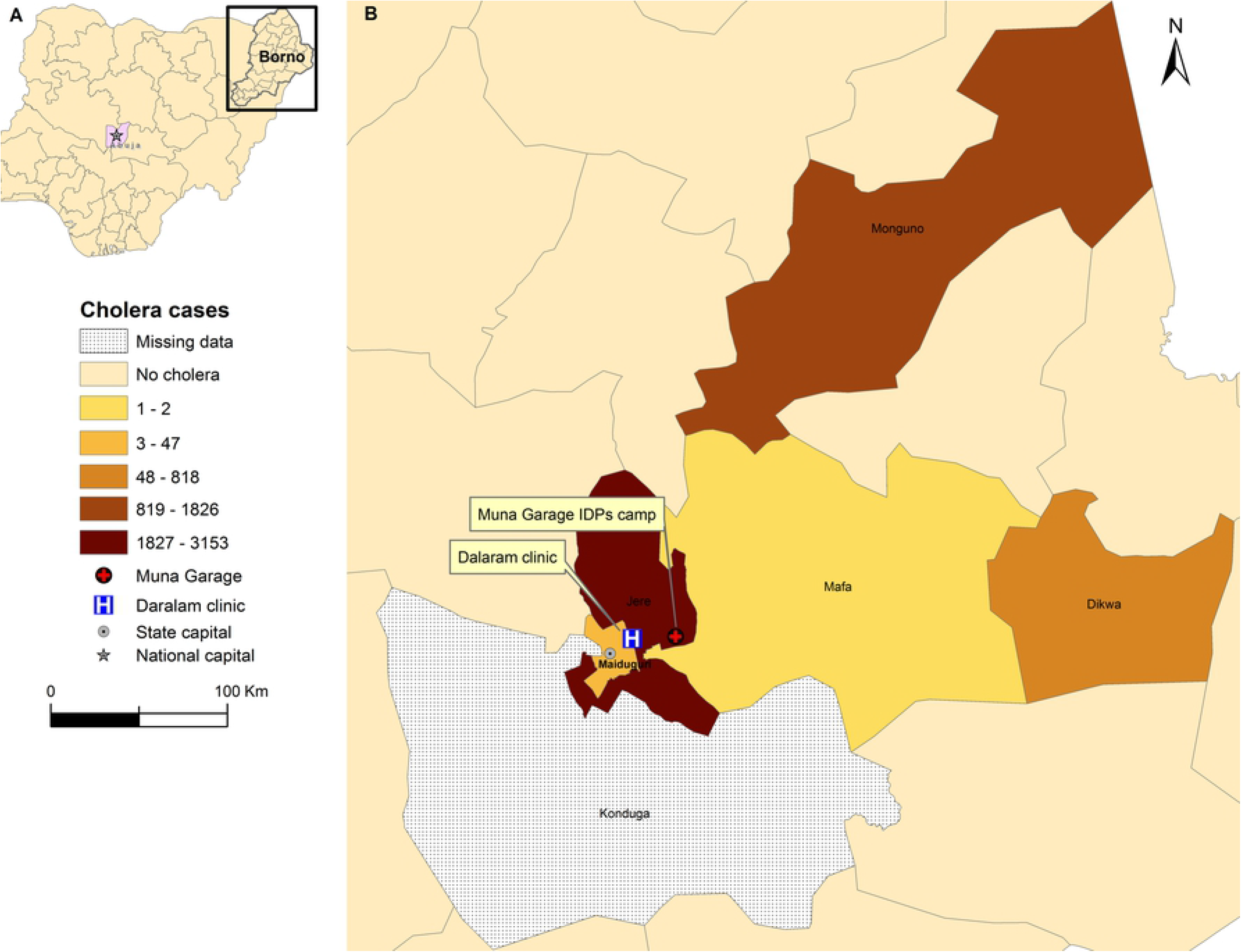
Spatial occurrence of cholera in Borno State, 2017. Insert (A) show Nigeria with national capital Abuja and Borno State, and (B) portrays the six Local Government Areas (LGAs) affected by cholera in 2017. The index case was detected in Muna Garage Internally Displaced Persons (IDPs) camp and the first cases where transported by ambulance to Dalaram clinic for treatment (insert B). Shapefile for Nigeria was obtained from WHO Nigeria Country Office as part of document review.

**Fig 2.**
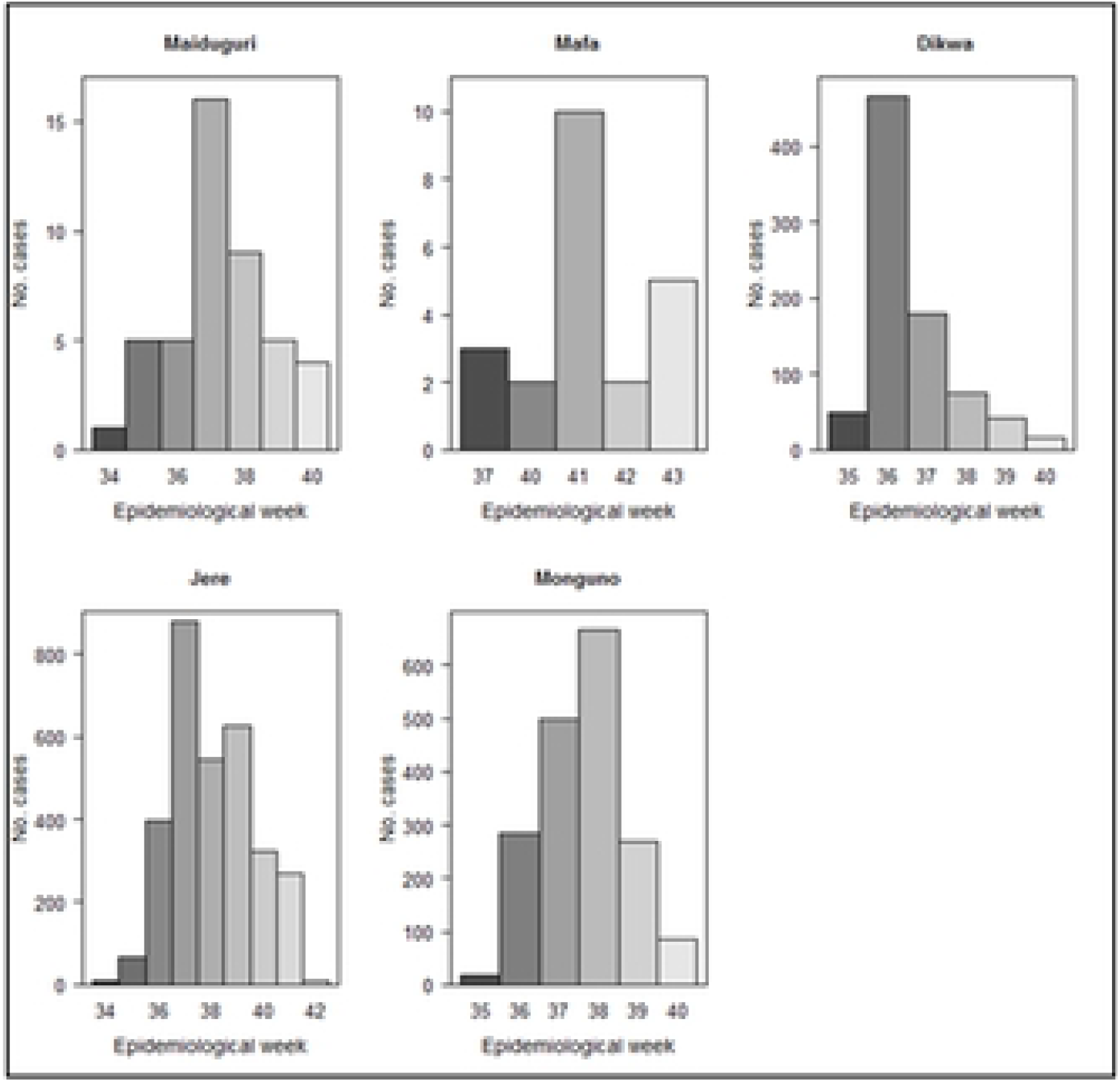
Weekly occurrence of cholera in Borno State, 2017. Data were not available for Konduga. There were no suspected cases of cholera in Borno State prior to week 34 and after week 42 of 2017.

### Evaluation of the Borno State cholera outbreak response

The overall response to the 2017 cholera outbreak in Borno State involved three main sectors namely; health, WASH (water, sanitation, and hygiene), and nutrition. With the inclusion of OCV, the main objective of this evaluation was to answer the questions: *What are the perspectives of government and emergency agencies such as United Nations (UN) and nongovernmental organizations (NGOs,) and how did partners coordinate an integrated response* which included epidemiology and laboratory surveillance, case management, OCV, WASH, nutrition, and cross-cutting issues such as coordination, communications, and logistics.

## Methods

### Study Setting

The qualitative study was conducted in Abuja, the capital of Nigeria, Maiduguri, the capital of and a Local Government Area (LGA) in Borno State, and Jere, a neighboring LGA to Maiduguri. Maiduguri and Jere are both urban LGAs in Borno. These locations were chosen because operational and administrative offices of government and partner organizations that provided interventions to the Borno cholera outbreak are located there. We conducted interviews and discussions among WHO Country Office, NCDC, Federal Ministry of Water Resources (FMWR), and NPHCDA emergency response field staff in Abuja. In Borno, the interviews and discussions were carried out amongst field staff and in the settings of WHO and UNICEF Borno State offices in Maiduguri, and Emergency Operation Centre (EOC) in Jere.

### Study design

An exploratory cross-sectional qualitative study design was used to gain experiences of the outbreak emergency response including timeliness and coordination of as well as constraints to effective response.

### Sampling

#### Selection of respondents

Participants were recruited using purposive and snowball sampling strategies [5]. Key informants (KIs) and group discussion participants were required to be representatives of organizations (governmental, international NGO, and United Nations (UN)) who took active part in the outbreak response. On the one hand, KIs were participants without location and time flexibility such as top ranking officials at MOH, WHO, etc. On the other hand, group discussion participants ranged from 3-7 members who had location and time flexibility for group discussions. At the national level in Abuja, a purposeful sample of participants were initially selected from WHO Country Office and a snowball strategy was applied to constitute subsequent organizations including NCDC, FMWR, and NPHCDA. At the state level in Borno, participants were initially selected from WHO Borno State office in Maiduguri and then extended outward to include respondents from BMOH, UNICEF, MSF (Médecins Sans Frontières), etc. A list with names, emails and telephone numbers of partner organizations that took active part in outbreak response was obtained from the WHO Borno office. Depending on the proximity of the partner to the WHO Borno office, participants were then recruited either in person, by email or phone. Subsequently, the recruited participants referred us to other partner organizations, who, in turn, recommended further contacts. The recruitment ended when participants could no longer recommend any new organizations. MOH health officials were interviewed at the EOC in Jere.

### Data collection

Data were collected by a team of two members, one interviewer (PhD researcher with training and experience in qualitative methods) and one assistant (note taker) with a Higher National Diploma in public administration and training in field note taking techniques, but without formal training in qualitative methods. The research was conducted from February 19-24 and 26-28, 2018 in English in Maiduguri, Jere, and Abuja (Fig 1A), respectively. The interviewer led the interviews, moderated the group discussions and audio typed the conversation(s) using a SONY audio (voice) recorder while the assistant took written notes as the participant(s) spoke. In interviews and group discussions where participants elected to be anonymous, only written notes were taken. Qualitative data collection methods incorporating key informant interviews (KIIs) and focus group discussions (FGDs) were used to understand the perspectives of both government and partners on the overall outbreak response. We used an interview guide and open-ended questions (S3 Supplement) to get the perspectives about the emergency outbreak response in both KIIs and FGDs. During FGDs, participants were encouraged to expresses their views in their own vocabulary, and explore further themes of interest to them. The duration of KIIs ranged from thirty to forty-five minutes and from one to two hours for FGDs. The data were encrypted and stored at Johns Hopkins University in a secure database for analysis.

### Data analyses

Tape-recorded interviews, discussions, and field notes were transcribed into English. In the transcriptions, important aspects of data interpretation such as voice speed, tone, and points of stress were captured. The transcripts were arranged according to government and partner data. The qualitative data analysis was inductive to ensure that the process focused on data. Focus groups and individual interviews were sorted and coded hierarchically into thematic areas. Data coding were performed using pen and paper. Computer software was not used. Some partners fit into more than one thematic area. In working through the transcripts, sections of text from KIs and FGDs that related to a thematic area were brought together. This was to ensure that the sources of the various texts could be re-traced. Thus, thematic similarities and dissimilarities of perspectives of the overall emergency response to halt the spread of the outbreak could easily be inferred.

## Ethical considerations

This qualitative study was conducted under the purview of WHO, NCDC, and BMOH to evaluate post outbreak response activities as part of monitoring and evaluation to inform future outbreak response efforts. Based on this, the Johns Hopkins University Internal Review Board (IRB) determined that the proposed study activity does not qualify as human subjects research as defined by United States Department of Health and Human Services regulations 45 CFR 46.102, and does not require IRB oversight. Further, as the study did not seek personal identifiers (name, birthdate, etc.) and relatively non-intrusive, verbal informed consent was obtained prior to conducting any interviews/discussions and after explaining the purposes of the study. To ensure the privacy of information, only authorized persons have access to the voice records and the list that indicates which records are associated with which respondents.

## Results/findings

### Thematic areas/pillars of the outbreak response

The purpose of this assessment was to understand individuals’ or groups’ perspectives about cholera interventions intended to stop the 2017 cholera outbreak in Borno State. During the outbreak, several organizations supported the national and State governments in the emergency response. The detailed evaluation included perspectives from 17 government and 22 partner representatives using KIIs, FGDs, and document reviews. Perspectives emerging from the data were grouped along three thematic sectors namely health, WASH, and cross-cutting. Analytic findings within these sectors fell into the following thematic areas namely 1. Health: epidemiology and laboratory surveillance, case management, and vaccination; 2. WASH: water, sanitation and hygiene; and 3. Cross-cutting themes including risk communication, coordination, and logistics.

### Epidemiology and laboratory surveillance

#### Outbreak detection and notification/reporting

Eight members in the surveillance team led by a State Epidemiologist were supported by WHO and other partners. Three platforms were available for epidemiological surveillance response, namely the Early Warning Alert and Response System (EWARS) [6, 7], Integrated Disease Surveillance and Response (IDSR) strategy [8, 9], and phone platform (alerts by mobile phones). Although the EWARS and IDSR were intended to quickly identify a cholera outbreak, there were different channels for communicating the cases during the early stages. In addition, there were discrepancies regarding the detection and reporting of the outbreak. From the WHO Surveillance Team in Maiduguri, the index case of the outbreak was detected and reported through the phone platform by MSF health facility in Gwange on Wednesday August 16th, 2017 as illustrated:

*“On August 16th, 2017, we got a call from MSF telling us about a case of acute watery diarrhea, which prompted an immediate visit to their facility in Gwange”* (KI, Dr. Uzoma Ugochukwu, WHO Surveillance Unit, Maiduguri).

But UNICEF also detected and reported another case probably different from the one reported by MSF.

*“A case was admitted in a supportive health care facility managed by UNICEF in Muna Garage and UNICEF-Health got the alert notification on Friday August 18, 2017”* (KI, Dr. Asta, Kone, UNICEF-Health Team Lead, Maiduguri).

*“Given that we were notified during an ongoing meeting and that it was very late that day and because of security reasons, UNICEF-Health couldn’t visit the health facility immediately. At about 8:00 am the following day, UNICEF-health visited the health facility in Muna Garage, verified, and confirmed clinical signs and symptoms of cholera from the household where the patients were coming. Upon confirming the case to be cholera, I immediately notified Dr. Izabelle Devaux, the then WHO Surveillance Team Lead, and the BMOH”*, Dr. Kone added.

In the disagreements as to which organization detected and notified the index case, we note that MSF detected and notified her case on August 16 while UNICEF detected and notified hers on August 18, 2017. Intriguingly, the two cases all originated from Muna Garage IDPs camp (Fig 1B).

On the government side, BMOH agreed to have received an alert from MSF about the outbreak,

*“The outbreak started on a site supported by MSF and they ‘<u>alerted</u>’ my office through the health sector coordination structure staffed with epidemiologists, disease surveillance and notification officers at the community, health facility, LGA, and state levels”* (KI, Dr. Lawi Meshelia, BMOH, Director of Public Health and Head of EOC, Jere).

The above excerpt indicates that MSF ‘alerted’ BMOH through the health sector coordination structure, and the word ‘alert’ signals that there was timely detection and notification of the outbreak. The use of the phrase ‘health sector coordination structure’ indicates that the phone platform was used to alert the ministry. At the federal level, the outbreak was reported to the NCDC through routine surveillance by BMOH:

*“We were notified through the routine surveillance system, as we saw an increase in the number of cases of cholera. The moment we saw reports stating cases were coming out from IDP camps, we decided not to take chances”* (FGD, Dr. Adesola Yinka-Ogunleye, Epidemiologist and NCDC Cholera Technical Working Group Lead, Abuja).

In surveillance, the word ‘alert’ portrays emergency and urgency to respond, but the word “notify” was used more frequently. What is mentioned above as routine surveillance refers to IDSR strategy [10], which was not directly mentioned during interviews. Furthermore, the Expanded Program on Immunization Team learnt about the outbreak while they were on vacation.

*“The cholera outbreak notification came in when we were on short break. We had to shorten our break and returned immediately to respond to the outbreak. Before our return, non-immunization activities had already been on ground, like WASH, active case search, and case management”* (KI, Dr. Oche James, Focal Person for Supplementary Immunization Activity, Maiduguri).

Meanwhile at the WHO Country Office level, the disease was identified through the weekly epidemiological reports of EWARS.

*“The moment the Office was notified through EWARS, it contacted the government (NCDC and BMOH) to make sure they declare it as quickly as possible”* (KII, Dr. Alemu Wondimagegnehu, WHO Country Office, Abuja).

Some partners specifically mentioned EWARS and phone platform surveillance systems, which are appropriate for reporting emergency that requires an immediate response. However, the IDSR strategy, although intended for early detection and rapid response to outbreaks with epidemic potential, was not mentioned. Other partners such as ALIMA and FHI360 were notified during a joint health sector meeting held at EOC.

#### Outbreak confirmation and declaration

Very critical to the outbreak response was outbreak laboratory confirmation by culture and declaration. As such, the call on August 16, 2017 prompted an immediate visit by WHO Surveillance Team to MSF Dalaram Clinic (Fig 1B) to ascertain the case. Despite timely visit, collection of samples at bed side, and a positive Rapid Diagnostic Test (RDT), the confirmation was delayed.

*“…the call from MSF on August 16, 2017 prompted an immediate visit to their facility in Gwange. We saw the case, took samples, and conducted RDT in the facility, which turned out positive. Further, we took samples and sent to the University of Maiduguri Teaching Hospital (UMTH) for confirmation, which turned out negative to culture. But because we saw the case and the stool that was classical of cholera, we sent part of the samples to the reference laboratory in Lagos, which turned out positive to culture”* (KI, Dr. Uzoma Ugochukwu, WHO Surveillance Unit, Maiduguri).

With a positive culture results from Lagos, a cholera outbreak in Muna Garage IDP camp in Jere LGA was confirmed. Nearly all partners stated there was delay in the confirmation as captured by the following excerpt.

*“There was probably a little delay in confirming the outbreak. I can’t really say how many days it took them the moment the case was identified, but they started managing it accordingly. From this end, the moment we were notified we contacted the government to make sure they declare it as quickly as possible”* (KI, Dr. Wondimagegnehu Alemu, WHO Country Representative for Nigeria, Abuja).

Nonetheless, BMOH disagree there was delay in case confirmation, stating:

*“Yes, the outbreak was declared timely because the issue of diarrhea and vomiting is all year round in this part of the country. Diarrhea and vomiting per say is not something that will excite anybody. So when we had these cases, nobody took it serious. We thought it was just normal treatment of diarrhea and vomiting and so on. But when the cases were becoming alarming, and the first confirmation on the 26th August, and by 28 of August, when we discovered that it was confirmed to be cholera, we alerted the highest authority, the Honorable Commissioner of Health. We advised him to come out and declare so that we can have synergy among all the partners that are in the State. So he now came out and declared the State as having outbreak of cholera”* (KI, Dr. Ghuluze Mohammed, Director Emergency Medical Response and Humanitarian Services, BMOH, Jere).

From above excerpt, the outbreak was confirmed 10 days after identification of the index case. Closely linked with the issue of outbreak confirmation was the issue of outbreak declaration by the State. The above excerpt from Dr. Ghuluze indicates BMOH declared the outbreak in a timely manner, which he added:

*“We also had to take into cognizance the political situation and repercussions of declaring an outbreak. We are in democracy. Somebody will just think that you are trying to score political points by declaring cholera outbreak. We had to be convinced and then convince the government to see reason to declare an outbreak”* (KII, Dr. Ghuluze).

Yet, as the outbreak started on August 16th, was confirmed on the 26th, and officially declared on August 28th (12 days after), several partners stated that outbreak declaration was delayed. The following excerpts sum up partners’ perspectives of outbreak declaration.

*“…Although it was a difficult task, but with much persuasion, clinical prove, and consultation, the state government declared the outbreak”* (KI, WHO State Coordinator, Maiduguri). *“There was this reluctance from the BMOH to declare cholera outbreak despite clinical evidence of transmission” (*KIs, Dr. Martinez, WHO Lead Coordinator and Dr. Issa Malam, ALIMA, Maiduguri). *“From the reports and surveillance data that were coming in, we knew we didn’t have to wait for the outbreak to be declared”* (FGDs, NCDC Technical cholera Working Group Team Lead, Abuja). Some attributed delayed declaration to vacation. *“The outbreak started when people were on vacation and did not want to cut short their vacation”* (Anonymous source). Others attributed the delay to the hajj, the pilgrimage to Mecca that every Muslim is required to make at least once in life. *“Declaring the outbreak meant that Muslims could not travel to Mecca, as is the policy that people from an active cholera region cannot travel to Mecca”* (Anonymous). Still, others took issue with WHO official policy to declare cholera outbreak: “*The policy that insist on declaring cholera outbreak only after culture confirmation despite positive RDT and clinical evidence of transmission is ill advised in settings where laboratory capacity is rare. In such settings, positive RDT and clinical evidence of transmission should suffice to declare a cholera outbreak. Had this been the case, the 2017 outbreak in Borno would have been contained at the inception of the emergency”* (Anonymous).

It is highly likely that lack of laboratory capacity such as reagents and trained lab technicians at the UMTH lab contributed to the negative culture results that affected timely confirmation and declaration. Without the declaration, surging up capacity in intervention measures such as oral rehydration points (ORPs), cholera treatment units (CTUs), and cholera treatment centers (CTCs) besides others could not be carried out. The declaration of the outbreak 12 days after onset likely slowed outbreak response efforts and contributed to its rapid spread from Jere LGA (Muna Garage IDP camp) to six other LGA.

#### Outbreak verification, investigation, and spread

Having been informed of the outbreak, the BMOH emergency took verification actions as typified:

*“On learning of the outbreak, we took actions including informing the local disease surveillance officers, primary health care coordinators, LGAs, and immediately mobilizing Rapid Response Teams (RRTs) to the community for outbreak verification in collaboration with partners like WHO, MSF, and others”* (Dr. Lawi Meshelia, Director of Public Health at BMOH and Head of Public Health Emergency Operation Center, Jere).

Although the BMOH emergency office dispatch RRTs and active search began immediately, still after initial appearance in Muna Garage IDP camp of Jere LGA, it appeared in five other LGAs. During this study, it was not clear what the mandate of the RRTs were and how effective and decisive they were in containing the outbreak within the camp. Results of the RRTs outbreak verification and investigation were not clear either. In contrast, the WHO-led surveillance team investigated and traced the first suspected case back to the household in Muna Garage IDPs camp (Fig 1B). The team found that sewage leakage from one of the latrines was flowing to the index case’s household.

*“When we went to the index case’s house, we saw sewage that was leaking into his house and we notified the WASH sector”* (KII, Dr. Uzoma Ugochukwu, WHO Surveillance Unit; Maiduguri).

They also found that almost all children in the index case’s household had cholera, which led to active case search in and outside the camp.

*“Upon discovery of the leaking sewage and because of our close partnership with the state government, the WHO Surveillance immediately contacted the State-WASH sector namely RUWASSA (Rural Water Safety and Sanitation Agency) and Borno State Environmental Protection Agency (BOSEPA) for repairs. When we returned one week later to decontaminate the leaking facility, it was still unrepaired,”* Dr. Uzoma stated.

RUWASSA and BOSEPA probably underestimated the magnitude the situation. This investigation illustrates that the 2017 Muna Garage IDP camp cholera outbreak may have originated from a leaking latrine. Of course, when latrines fill-up and leak, cholera ensues.

Within a week cholera spread to the whole camp and host communities. Some key factors that may have led to the sporadic spread of the disease were delayed declaration and festive period of ‘Eid El Jabir, and the delayed establishment of the CTC at Muna Garage IDPs camp (Fig 1B).

*“The time lag between the period of waiting for culture confirmation … created room for the wide spread of the disease”* (KII, Dr. Uzoma Ugochukwu, WHO Surveillance Assistant Team Lead).

*“Immediately after Eid EL Jabir, cases were reported in other LGAs. Realizing the potential of Eid El Jabir in the spread the outbreak, awareness was raised in response, but restrictions on population movement could not be undertaken”,* Dr. Uzoma supplemented.

Echoing Dr. Uzoma, Dr. Okudo asserted,

*“The outbreak spread rapidly, because there was this festival with lots of eating and celebration and after which there was a jump in the number of cases. So we knew we had a full blown outbreak”* (Dr. Okudo Ifeanyi, WHO Acting TL&WCL, Abuja).

Eid El Jabir, a Muslim religious festival that entails lots of movement and food sharing and that coincided with the outbreak, may have amplified the outbreak. Furthermore, though the outbreak started in Muna Garage IDPs Camp of Jere LGA, the first CTC was set up in Dalaram Clinic in Maiduguri LGA (Fig 1B).

*“Basically, this disease spread out as a result of lack of CTCs in Muna Garage IDP Camp at the early phase of the outbreak. The response to the outbreak at the initial stage was handled too slow, thinking it could be managed with a Primary Health care facility. Unfortunately, no CTCs were set-up in Muna Garage IDP Camp. Rather, it was setup in Dataram Clinic, some few Km away from Muna Garage. As such, cholera patients were referred from Muna Garage to Dataram Clinic using ambulances, which exposed anyone in the ambulance to the disease. It was when the number of cases was going up and an advocacy made to the Commissioner of health that a decision was reached to set up a CTC within the Muna Garage Camp”* (Dr. Asta Kone, UNICEF Health Team Lead, Maiduguri).

Dr. Alemu adds more insight to Dr. Kone’s account vis-à-vis tardy CTC set up in Muna Garage camp:

*“The issue was disagreement between BMOH and partners in selecting the site for the second CTC. BMOH wanted it within the camp while the other partners wanted a little bit further. This definitely delayed the CTC construction”* (KII, Dr. Alemu, Wondimagegnehu, WHO Country Representative for Nigeria, Abuja).

Fig 1C shows the location of Muna Garage IDP Camp and Dalaram Clinic, a nutrition center in Maiduguri catering for malnourished children of the Boko Haram crisis. Transporting cholera patients from Muna to Dalaram Clinic resulted from disagreements and was risky for cholera transmission as this exposed uninfected persons with the danger of contracting the disease.

Delayed declaration (state needed positive culture to declare outbreak), lack of coordination between WASH and Surveillance teams in sharing actionable information, and disagreement between BMOH and partners contributed to a delayed response including WASH, case management, and behavioral change communication.

### WASH (water, sanitation, and hygiene) response

There were 12 organizations in the WASH team led by the RUWASSA with UNICEF-WASH playing a crucial role in the response activities. During the outbreak response, the NCDC Epidemiologist played the role of coordination and report presentation of WASH interventions.

UNICEF-WASH early outbreak investigation in Muna Garage, corroborated those of WHO-led Surveillance Team, documented several fecal oral transmission routes.

*“Our immediate outbreak investigation showed several transmission opportunities such as latrines fill-up and overflow, which were the top most risk factors of the outbreak; less than half a meter deep shallow latrines inside houses and tents; open defecation around refuse dumps, non-functional, and/or collapsed latrines; some families disliking communal latrines and others considering communal latrines very far from their tents; large stagnant pool of water draining from road side into the camp in which people washed cars, bath, and kids played; and overcrowding in the camp”* (KI, Mr. Kabuka Banta, UNICEF-WASH Specialist, Maiduguri).

The living conditions in the camp were highly favorable to cholera outbreak. Of course, if latrines collapsed or overflowed, families will dislike using them and turn to open defecation, a practice they are used to in the villages from where they fled. Changing practices of open defecation has been a challenge. Likewise complaints about distance of latrines to the houses/tents and having less than half a meter deep latrines in houses/tents are typical practical ways of circumventing inaccessible communal latrines. Overcrowding in the camp likely forced families to line up to relieve themselves using the limited number of latrines, and this lead to their collapse and eventually being non-functional. All these conditions were compounded by stagnant pools of water draining into the camp in which children played.

After UNICEF-WASH identified transmission routes, it mitigated these off by 1) providing aqua taps to households, 2) equipping about 650 volunteers with quick testers to test water contamination, 3) engaging street water vendors about water safety, 4) promoting good hygiene behaviors through social mobilization, 5) dislodging and chlorinating latrines, and 6) chlorinating water spots/tanks at household and community levels. However, the lack of first mapping, i.e., audience stratification by tribe, language, religion, and leadership, the IDP camps and host communities to understand the audience led to misconception of chlorine as a sterilizing agent. This led to avoidance of chlorinated water and latrines as well as increases in open defecation.

*“As technicians doing WASH we know what is required to curtail an outbreak, but we don’t know exactly what method to adapt. For instance, in our immediate response to the outbreak, we chlorinated water and latrines but these led to the avoidance of the chlorinated water and latrines as well as increases in open defecation”* (KII, Mr. Kabuka Banda, UNICEF-WASH Specialist, Maiduguri).

Without fully understanding the people’s reluctance to use chlorinated water and latrines and their preference towards open defecation, UNICEF-WASH engaged UNICEF Communication for Development (C4D).

*“As such we engaged C4D and in less than two days, they came up with tenable solution, which the WASH team failed to recognize. There were stratifications within the community based on their faith, language groups, and leadership, which WASH couldn’t initially identify. The introduction of the locals gave us a lot of feedback as to why there was reluctance to use chlorine. It was discovered that there was misconception about chlorination. Some were afraid that chlorine was for sterilization, which would stop women from giving birth if they use the chlorinated water. There was also this misconception that chlorinated latrines produce chlorine vapor and when women use them the vapor will sterilize and stop them from giving birth,”* Mr. Kabuka Banda continued.

We note here that the technical people did not use the phrase ‘sterilize water’, which the community could have mistakenly thought to mean sterilize for fertility. What was actually rumored in the community was,

*“You see that whitish think that they are putting in water, houses and latrines, it will stop women from giving birth to children” (KII, Dr. Gerida Birukila, UNICEF-C4D, Maiduguri).*

The identification of the community structure uncovered these misconceptions and so the question then become who amongst these strata do the people trust to talk about promoting chlorine and personal hygiene?

*“Working with C4D, the WASH team mobilized more than 430 people within the community who then became more effective in transmitting campaign messages and eventual chlorine acceptance. In other words, in order to have a successful intervention, it all boils down to who are involve”* (Mr. Kabuka Banda emphasized).

Key among the people that were mobilized in the community were ‘Bullemas’, community leaders and gate keepers in whom people trust and who have extensive power in community decision making. Attempts to provide interventions in community without consent of ‘Bullemas’ encountered resistance no matter how well intended the interventions were. C4D was successful because they sought first to understand community structure through ‘Bullema’ networks and door-to-door communication strategies before lunching behavioral change intervention. The networks were built under the polio eradication platform.

Similar to UNICEF-WASH in Muna Garage, Family Health International 360 (FHI360) emptied filled latrines in Dikwa LGA in response to the outbreak.

*“Where this outbreak started in Dikwa, we saw that 75% of the cases where coming from one place called ‘Bulabulin,’ where there was a filled latrine. This filled latrine was dislodged by FHI”* (FGD, Dr. Kibebu Kintu Barta, Primary Health Coordinator, FHI360).

Additional WASH responses in Dikwa included chlorination of water points, household water buckets, and latrines; distribution of Aqua-taps; dislodgment of filled latrines; community sensitization; and hygiene promotion awareness.

As concerns WASH response from the FMWR, the latter was not notified of the 2017 Borno State cholera outbreak and therefore was not involved in the outbreak response.

*“The Federal Ministry of Water Resources intervenes only at the request or when it gets report from the State Ministry of Water Resources of a need for intervention, which was not the case* (KII, Mr. Olu-Daniels Ibiyemi, Deputy Director of WASH at FMWR, Abuja).

*“But UNICEF did work with Borno State RUWASSA to provide WASH interventions during the outbreak,”* Mr. Olu-Daniels added.

Apparently the FMWR was not involved because she was not notified. Perhaps Borno RUWASSA did not see reason to involve the Federal ministry but WASH conditions in the camps prior the outbreaks indicate the need for more collaboration between Federal and State water ministries, and UNICEF.

*“There is need for UNICEF to support the FMWR, just as WHO does for NCDC, to carry out interventions and the need for the Ministries of Health and Water Resources to work together to combat water borne diseases,”* Mr. Olu-Daniels stressed.

The call for UNICEF to support the federal ministry signals that more collaboration is needed between government and partners in WASH. This need for collaborative support is clear when one looks at the WASH conditions in the camps prior to the outbreak as documented by various sources. Collaboration is also needed between federal and state water ministries given that the former played no role in the outbreak response. The following were challenges encountered by the WASH teams: 1) misconception about chlorine (it was associated with sterilization) led to avoidance of chlorinated water and latrines, 2) at peak season electricity from solar panels was not enough to provide water into the communal thanks, so people turned to street vendors, 3) unemptied, collapsed, and leaking public latrines compounded open defecation, a behavior the people were conversant with.

### Case management

Case management teams consisted of nine members led by the Borno State Hospital Management Board with NCDC playing a coordination role and partners such as UNICEF-Health, MSF, Medicins Du Monde, ALIMA, and FHI360 running health facilities in various LGAs. Depending on the state of dehydration patients were treated either in one or all three type(s) of facility namely ORP or CTU or CTC, which made it hard for locals to decide which to visit as exemplified.

*“About two days later, MSF set up an ORP clinic close to the UNICEF CTC which became conflicting as the locals don’t know the difference between acute watery diarrhea and cholera”* (KII, Dr. Asta Kone, UNICEF Health Team Lead, Maiduguri).

The setting up of the MSF ORP, though with the best interest of patients in mind, presented confusion for patients about where to seek care. It also added extra effort from the ministry to coordinate since it was also running a CTC in Muna Garage IDP camp with support from WHO.

*“Our CTC had 135 workers including 70 health workers, five doctors, 15 nurses besides 45 community health and extension workers and sanitation van guards. The State also provided the police, security, civil defense and military at that center and paid all stipends throughout the response”* (KII, BMOH, Jere).

In camps where the three facilities were operated by three different partners, issues of competition arose as summed up in the excerpt of one of the partners.

*“The only challenge we can clearly point out was the unnecessary competition amongst partners over who takes the glory?”* (FGDs, FHI360, Maiduguri).

This competition manifested in many ways including case referrals as exemplified.

*“MSF France provided case management in Jere in the Muna Garage IDP camp but referred cases to CTC ran by MSF Belgium in Dala IDP camp, where the later provided case management”* (KII, MSF Spain, Abuja).

As such patients who visited MSF Frances’ ORP in Muna Garage camp where transferred to MSF Belgium’s CTC in Dala, some kilometers away, because she was not permitted to set up a CTC in Muna Garage. Yet, there were two CTCs in Muna Garage, one run by BMOH and the other by UNICEF-Health. Therefore, it is counter intuitive that cholera patients from Muna Garage were referred to Dala.

Still in light of patient referrals, equally challenging were patient reluctance/refusal of transfers to other camps and health facilities, particularly, when they perceived disagreements between partners. However, FHI360 reported no issues in referrals as they had an integrated case management setup.

*“In Dikwa, where we provided case management in a government-owned but FHI360-run health facilities, our health outreach workers referred patients with acute watery diarrhea directly to ORPs managed by FHI360 and from there to CTUs or CTCs equally managed by FHI360…our strength was not only this in-house referral system but also our willingness to work with other partners”* (FGDs, FHI360, Maiduguri).

Throughout this study, we documented that challenges with referrals were more common in inter-partner transfers and minimal in intra-partner situations. Other issues mentioned that affected case management response were late setup of the CTCs in the Muna Garage camp, denial of cholera by the locals, shortage of trained and experienced personnel to deal with emergency, and underestimating the threat the outbreak posed.

### Oral cholera vaccination

The vaccination campaign team had five members under the leadership of the State Primary Health Care Development Agency (SPHCDA) with support from NPHCDA including partners such as MSF, UNICEF and WHO Health Operations Team. This thematic pillar also included 27 Disease Surveillance and Notification Officers (DSNO) at the LGA level. Remarkably, this team planned, applied for, and deployed OCVs in Borno State within two weeks; Understanding how these were achieved was one of the main objectives of this study.

#### Vaccine application and Micro Planning

The use of oral cholera vaccines in Nigeria was first considered in the context of endemic use as part of national cholera preparedness plan for 2017 during a workshop organized by NCDC in Abuja between May 31^st^ and June 1^st^, 2017. The idea of using OCV came up during this preparedness workshop as exemplified:

*“Between the 31st May and June 1st, 2017, NCDC had a cholera preparedness workshop where states with history of cholera epidemics were invited including Borno State. Partners were also invited to the preparedness workshop including representatives from the WHO headquarters. It was at this workshop that the idea of using OCV in Nigeria was first discussed … and immediately NCDC wrote to the various partners and stakeholders seeking their consent to use OCV. The outbreak in Borno now gave the opportunity to use OCV in the country for the very first time. It is important to note that the cholera preparedness workshop was not in response to the outbreak in Borno”* (FGD, Dr. Adesola Yinka-Ogunleye, Epidemiologist and NCDC Cholera Technical Working Group Lead, Abuja).

The 2017 Abuja preparedness workshop laid the foundation for OCV use in Nigeria. One of the recommendations from the workshop was the approval of OCV in Nigeria, which NCDC applied and obtained the approval from the National Agency for Food and Drug Administration. Thus, the August 2017 Borno outbreak gave opportunity to use the newly approved intervention, and in so doing turned endemic consideration quickly into reactive (emergency) use with initial focus in Muna Garage IDP Camp (Fig 1B). Key to the deployment of OCV was the existing robust polio vaccination structure:

*“The most important success factor of the OCV response from the planning to operational phases was the implementation using polio structures, an existing, robust, highly established, and functional structure, which made the OCV response easier and swifter. For example, micro planning that was used in Sierra Leon was obtained and with simple adaptations of polio mechanism and assistance from Headquarters, Micro Planning for Borno was ready and OCV request submitted within days. WHO Surveillance Unit provided data that was used in the Micro Planning and OCV application”* (KI, Dr. Dereje Ayana, WHO Health Operations Team Lead, Maiduguri).

In line with above excerpt, WHO Country Office underscored the role of the polio structure, which facilitated rapid importation of OCV into Nigeria and in the field in Borno within two weeks:

*“With regard to the vaccination … Firstly, we (government, WHO, GAVI, UNICEF) managed to bring the cholera vaccines in two weeks. Within three days the application was submitted. The next week, it was approved and the next five to six days it came to the country and were shipped to the affected region. Secondly, the polio mechanism, with its good practice and experienced staff, was used for the OCV, and so it was easy to have OCV without having to go through training vaccinators and stresses including logistic support. Thirdly, the Lot Quality Assurance Survey helped us to know the degree of its quality. In addition, advocacy from the state was very high including the Commissioner vaccinating to show that OCV is harmless, and that it protects. Advocacy was well done, everybody was involved, finance, and the vaccines were secured from “Global Alliance for Vaccination and Immunization-GAVI”. Lastly, Well-coordinated structure under the leadership of the EOC management, and experienced workforce that have the experience conducting immunization…combined together and made it very easy for us to achieve a remarkable accomplishment over the OCV”* (KII, Dr. Wondimagegnehu Alemu, WHO Country Representative for Nigeria, Abuja).

The most tedious part of any vaccination campaign is developing the Micro Plan because it needs the population structure of target population, which is often lacking, to estimate vaccine numbers. But with population data from WHO Surveillance Team and OCV Micro Planning guides for OCV campaign from Sierra Leone, Micro Planning for Borno was ready within days. The polio team has an existing and flexible structure for rapid implementation of immunization campaigns, and as such, polio personnel and other partners were involved from the micro planning stages leading up to the field implementation.

Adaption of the polio platform for OCV needed training to accommodate OCV unique peculiarities, which was done at various levels (S1 Fig):

*“Accordingly, polio staff were trained at the national, state, LGA (DSNOs), ward levels, and down to field polio vaccinators to handle OCV* (KII Dr. Dereje Ayana, WHO Health Operations Team Lead, Maiduguri).

Notwithstanding extensive polio staff training (S1 Fig), there were still challenges in OCV administration. First, the vaccination campaign team had to work extra hard to convince the adult population that they needed OCV. This demographic group associated vaccination with children. Polio vaccinators were not familiar with opening OCV vials and so were provided scissors to overcome this difficulty. Vaccination cards were not issued during the first round of vaccination, with the assumption that there will be no second round. Although the latter subsequently became apparent efforts during the second round to document reception of vaccine during first round proved futile. Data flow from LGAs to the central coordination with poor communication network were slow and created anxiety as to whether relevant data would be available to inform needed subsequent actions. To overcome this, supervisors travelled to the villages to get the relevant data for action. In some instances, partners put self-recognition over public health interest asking who gets the glory. Who owns the data? However, these faded when partner’s attention were called to the fact that all data belongs to the ministry. In an attempt to save time, plans to transport vaccines from Abuja to Borno by road were abandoned in favor of flight transport, as timeliness of the vaccine delivery in Borno was critical. Limited cold room capacity in Abuja and Borno led to vaccines being transported in batches, which was not very cost efficient.

#### Vaccine finance

The existing polio structure blueprint was used to model the OCV budget because Borno State has a strong polio platform. For the budget preparation, four working groups were formed (Technical, Training, Social Mobilization, and Logistic Working Groups) saddled with the responsibility of drafting a budget using the polio blueprint. These drafts were merged, submitted to, and approved by GAVI, and from there, the funds were sent through the WHO Country Office to the local WHO Health Operations Unit. This Unit then paid both government and partner beneficiaries associated with the campaign.

Although the existing polio structure payment system was adapted, its card mode of payment (S2 Fig), which handles situation where there are no banks/bank accounts, was not adapted:

*“The WHO funding system emphasis direct disbursement into beneficiaries’ bank account, which was frictionless where beneficiaries had bank accounts, i.e., up till the LGA level”. This became problematic at the ward level where people lack access to bank let alone having a bank account. In this situation, ward level people were paid cash through the Local Immunization Officer, who, in turn, retired back to the WHO Finance Unit. The state LGA and WHO partner cluster coordinators monitored the whole payment processes to ensure all participants were paid appropriately”* (KII, Finance Officer, WHO Health Operations Unit, Maiduguri).

The polio card mode of payment could also be adopted in case of emergencies like the OCV. The polio card system entails contracting with credible financial institutions in the country to pay beneficiaries upon presentation of the cards. This discourages physical cash payment and saves the finance unit the stress of going to the field to make payments. The major challenge to the finance team was lack of bank accounts at the ward level. To overcome this, the LIO’s or WHO ward focal person’s account was credited to make physical cash payments. In addition, the finance team under-budgeted during the first round, which posed a threat to the success of the campaign in some LGAs. To make up for this mistake, team members had to work extra days with less pay. This mistake was corrected during second round budgeting.

### Cross-cutting themes

#### Risk communications

The Communications team was made up of six organizations led by the State Primary Health Care Development Agency (SPHCDA) with the NPHCDA, WHO, UNICEF, and other partners playing critical supportive roles. Social mobilization was under the prevue of WHO, UNICEF, BMOH and other organizations. WHO communication response was a three pronged approach encompassing 1) risk communication, 2) advocacy/visibility, and 3) raising awareness.

*“We created Outside Broadcasting System, which used speakers to communicate cholera risk messages, as most of the affected community did not have access to mass media such as television, radio, and power supply while advocacy and visibility engaged mass media channels to project WHO’s work to the entire world. Further, awareness raising distributed flyers house-to-house addressing Frequently Asked Questions, basic facts about cholera and vital information prevention”* (KII, Dr. Chima Emmanuel, WHO communications, Maiduguri).

About 25 mass media channels were engaged and taken to the field to report WHO staff doing surveillance, mapping and taking samples. These media outlets included four international radio stations, one international TV, and ten newspaper outlets. Besides publishing cholera stories on the WHO website, the communication team also created pictorial awareness campaigns using different local languages and passed these across to the different IDP camps across the state to raise awareness. The communication team also engaged a local theater and drama art group that staged plays in various IDP camps, which yield some positive results with positive behavioral and attitudinal change. Geo-coordinate maps guided WHO communications team in prioritizing and effectively delivering messages.

The WHO communications team used a top down approach to map and understand the audience. However, the first communication response to an outbreak should be mapping IDP camp and community structure by communication experts to inform response strategies. For instance, initially, cholera awareness raising did not ensure full participation of the community because key messages were in Hausa as mobilisers assumed Hausa was spoken widely. The key messages from the language changed to ‘Kanuri’ when communications experts mapped camps and found ‘Kanuri’ was the predominantly spoken language.

Meanwhile, UNICEF-communication for development (C4D), the branch responsible for communications within UNICEF, used a bottom up approach to first understand community structure/stratification through ‘Bullema’ networks and door-to-door communications before lunching behavioral change interventions.

*“The results of our one-on-one communication in the IDP camps indicated that less than 10 people in the camps had radio, that about 12 tribes lived in the camps, and that Hausa was not predominantly spoken as was initially assumed. Thus, our social mobilization was then based in the latter findings, which yielded an understanding of not only the channels and languages of communication but also the content of the messages”* (KII, Dr. Gerida Birukila, UNICEF-C4D, Maiduguri).

Of course UNICEF-C4D was already working in the community under the polio eradication platform before the outbreak and so had a network of people. Using the bottom up approach, UNICEF-C4D stratified the community by tribe, religion, leadership, and languages leading to a change in the main language of communication from Hausa to Kanuri. In addition, channels of communication changed leading to the appreciation of the initial reluctance and skepticism about chlorine, which UNICEF-WASH ignored. This was compounded by deep mistrust of information from government owned and run media.

### Logistics

The logistics team was under the leadership of BMOH buttressed by National Primary Health Care Development Agency, and WHO-OSL (Operations Support and Logistics). WHO-OSL was tasked with several roles including operationalizing the 2017 emergency preparedness plans, setting up CTCs, arranging/mapping inventory, and OCV logistics.

*“The 2017 cholera preparedness plan happened in May/June before rainy season, and WHO-OLS prepositioned 22 cholera kits in strategic locations in Borno, according to the preparedness plan”* (KII, Dr. Muhammed Shahid, WHO-OSL Team Lead, NE, Maiduguri).

When a decision was taken to set up a CTC in Muna garage, WHO-OSL had just four days to set up the CTC with its critical zones and to train staff for its proper usage to avoid contamination during treatment.

*“In setting up CTCs in Muna Garage, this was needed within less than 4 days, which was timely setup to include 1) neutral entrance zone for staff and 2) patient entrance zone (patient entrance, registration, out/in-patient (male/female), surveillance, recovery, waste management, deaths, and exit). Also WHO-OLS trained staff, supervised waste management, provided packaging materials and sample transport, and maintained cool chain. In addition, WHO-OLS mapped cholera supplies between BMOH and partners (e.g., FHI360-Dikawa, MSF-Monguno) according to availability (who has Ringer’s lactate? Tents? Kits?), and capacity to respond”* (Dr. Shahid noted).

WHO-OSL faced challenges taking inventory of partner supplies to avoid shortages as some partners were hesitant to share their stock while others lacked data. Yet mapping supplies did uncover critical shortages in Ringer’s lactate, which could not be purchased in large quantities in Borno, prompting WHO-OSL to act assertively to import the product.

*“As purchasing Ringer’s lactate locally was not allowed, WHO-OSL networked with WHO Headquarters, Regional Office and with MSF-validated supplier and 40,000 liters of the product was imported on time”* (Dr. Shahid explained).

It was not clear why Ringer’s lactate could not be sourced locally. As the second phase of response focused on OCV, WHO-OSL participated in Micro Planning design and cold chain maintenance in terms of making sure ice packs, vaccine carriers, refrigerators, and cold room storage besides freezing capacity were available.

*“As OCV used the **polio structure**, which is very strong in Borno, OCV logistics went smoothly. However, polio cold chain was limited and so work-around strategies such as bringing the vaccine to Maiduguri in phases using chartered planes were adapted. The real challenge was moving the vaccines to LGAs as this required military escorts* (Dr. Shahid said).

At the port of entry, National Strategic Cold Store (NSCS) under NPHCDA received the vaccines in Abuja and immediately shipped to Borno for lack of storage.

*“We received and immediately moved the vaccines to Borno State through trans-docking because of the storage constraints in Abuja warehouse and because of the policy in the country regarding supplementary vaccines. At every stage of the transport NSCS updated relevant stakeholders including NCDC, BMOH, and WHO”* (FGD, Mr. Palman USMAN, Head of Operations, NSCS-NPHCDA & Ms. Hauwa N. Tense, Warehouse Manager, NSCS-NPHCDA, Abuja).

The most essential action for the vaccine importation was a joint planning meeting where the roles and responsibilities of every partner were clearly defined. For timely stakeholder update, a pre-alert plan was designed to speed up the movement of the vaccines to Borno. This was in close collaboration between NSCS, BMOH, WHO, manufacturer, and the transport agents. When the vaccines arrived, NSCS sorted them out based on the batch no, expiry date, and carried out sampling following the normal store procedures. Due to storage constrain in Borno, vaccines arrived in batches for ease of deployment to LGAs.

### Coordination

The coordination thematic pillar headed by BMOH consisted of five members from the Federal MOH, NCDC buttressed by nine members from WHO, two from UNICEF, one from MSF, and one consultant. The BMOH emergency coordination office took the lead in coordinating partner activities including arranging military escorts for partners who could not afford their own security.

*“As health sector coordinating office, my role was tied with coordinating all the activities of partners from surveillance, WASH, RRTs, to setting up CTCs and to management of the centers besides coordinating military escorts for all health teams by liaising with the military state commander in charge of security”* (KII, Dr. Mohammed Ghuluze, Director of Emergency Medical Response and Humanitarian Services, BMOH, Jere).

The director further detailed what coordinating the overall response meant in terms of avoiding duplication of efforts and ensuring data harmonization.

*“Coordination meant coordinating partners such as WHO, MSF, FHI360, UNICEF, ALIMA, etc. and sending them where their services where needed in the different LGAs and IDP camps and to ensure there was no duplication of effort. It also meant ensuring that in LGAs where two or more partners worked, data coming out were harmonized; only one data coming out from the LGA, not data from UNICEF or WHO or MSF… As a coordinating mechanism, disease control officers and data managers from human machine interface (HMI) office were constantly collecting data from partners, which were used for planning, coordinating, and decision making”* (Dr. Ghuluze affirmed).

Double counting was at issue in IDP camps in which two or more partners worked. The ensuing excerpt illustrates why data coordination to ensure harmonization in a LGA was critical.

*“There were challenges in counting patients. Some patients visited all three health facilities moving from ORP to CTU to CTC operated by different partners. In reporting, ORP partner reported say 12 patients, then CTC partner reported say 14 patients etc. But a close review showed patients have gone from an ORP to a CTU and to a CTC. The partners, justifiably, needed to show they did work and this resulted in double counting. As such partners were allowed to count their respective contributions for their reporting and accountability and not the number of patients”* (KI, Dr. Okudo Ifeanyi, WHO Acting TL&WCL, Abuja).

The constant collection of data from partners by HMI officers was designed to overcome double counting and partners’ claim over case data. Yet, at the time of this research, data were being awaited from some international NGOs. Meanwhile, coordinating budget between government and partners at the early phase of outbreak response was not readily apparent.

*“Coordination of budget and funding between government and partners was not clear at the start. It was difficult to get what was needed for the response in terms of what was available with partners and what the government needed to contribute”* (KII, Dr. M Lawi, Director of Public Health, BMOH & Head of EOC, Jere).

For an effective response, the coordination team needed to overcome not only hurdles in budget allocation between partner and government, but also hesitancy in team formation. Some partners were reluctant to be part of the team. Other issues affecting coordination at the <u>start</u> of the response included some partners not coming in through a single identified channel, competing demands between insecurity and health emergencies, and the lack of synergy between the WASH and health sectors.

*“WASH was doing its own activities and so was WHO,”* Dr. Lawi asserted.

The mention of WASH above refers to UNICEF. As the outbreak ranged on, the need to overcome coordination difficulties became dire. In this respect, the completion of the EOC by WHO and handing it over to the BMOH just at the onset of the outbreak could not have happened at a better time.

*“The greatest success in overcoming coordination hurdles was the EOC, which was used as the planning and response center for all partners. The second factor in coordination break through was the State identification of the key thematic areas and populating these with key people in teams of between 5 and 7 members,”* Dr. Lawi continued.

NCDC agrees, emphasizing the role of EOC in coordination break through.

*“In a bid to respond to the outbreak, our main focus was to work with the state and partners to coordinate response. It was about that period that the EOC was setup by WHO and just commenced operations and enabled a meeting point between all partners”* (FGD, Mr. Womi-Eteng, NCDC, Incident Coordination Center Team Lead, Abuja).

In this vein, NCDC used EOC to form the Incident Management Structure (IMS), which consisted of an Incident Manager, the different pillars, and team leaders, to facilitate coordination. EOC made possible synchronization amongst thematic pillars including WASH, surveillance, logistics, case management, laboratory, social mobilization, vaccination, and risk communication. Coordination meetings where held at EOC at <u>4pm daily</u>.

*“Until declaring the State free from cholera, there was joint government and partner meeting at the EOC at 4pm daily where every team lead debriefed other team members of the situation on the field and data clearly analyzed to ascertain progress made. In areas and communities without network, data was sent using Thuraya or Satellite Phone”* (KII, Dr. Dereje Ayana, WHO Health Operations Team Lead, Maiduguri).

During the 4pm daily meetings, epidemiological curves and geo-coordinate maps were presented to direct multi-sectorial teams regarding which locations (recently affected households and surroundings) should be given priority. The use of Thuraya satellite phones to get data from network inaccessible locations highlights the importance of making data driven decisions.

Dr. Jorge Martinez lauded the EOC in facilitating and galvanizing coordination of the overall response efforts when he remarked,

*“EOC enabled the BMOH to take leadership and ownership in coordinating the outbreak response”* (KII, Dr. Jorge Martinez, WHO Health Sector Coordinator, Maiduguri).

*“On the other hand, coordination of response could have been better had WASH on the ministry side, i.e., RUWASSA, in terms of leadership of the sector, did not rise to the occasion with a slow start,”* Dr. Martinez added.

WHO State Coordinator, Dr. AM Taiwo, agreed with the previous remark about EOC and RUWASSA when he noted,

*“The setting up of the EOC by WHO was a blessing because it enabled coordination between Expanded Program on Immunization team and all emergency teams in Borno to carry out joint activities. However, WASH response was not as prompt as expected”* (KII, Dr. Audu Musa Taiwo, WHO State Coordinator, Maiduguri).

The late response from RUWASSA in taking leadership in coordination of WASH interventions ties with previous insights on WASH conditions in Muna Garage IDP where the outbreak started. Further, perspective from the FMWR indicated they were not informed of the outbreak. Had RUWASSA notified the FMWR of the outbreak, they may have had support from them just as BMOH had from NCDC. It was a missed opportunity. Overall, both partners and government concur that key to overcoming coordination hurdles were a functioning EOC, which enabled a meeting point amongst emergency thematic pillars, implementation of IMS, and inter-sectorial multi-agency debriefings at 4pm daily.

## Discussion

This qualitative research explored perspectives of reactive responses to stop the cholera outbreak emergency that started in the Muna Garage IDPs camp in Borno State, Nigeria, on August 16, 2017. The first cases were identified quickly by RDT but there was a considerable delay in declaring the outbreak because this depended on culture confirmation. We found that the epidemiological surveillance system (EWARS [6], IDSR [10], and the phone platform) quickly detected and notified health authorities of the outbreak. The system also classified the index case as fitting the WHO’s standard case definition and obtained a positive RDT test from the index case [11]. Nonetheless, laboratory confirmation of the diagnosis was a challenge. The samples with positive RDT result at bedside tested negative for *V.* cholerae using culture methods at the local UMTH laboratory; and thus, delayed outbreak confirmation as recommended by WHO [11]. Subsequently, *V.* cholerae was confirmed by culture in the reference lab in Lagos on August 26, 2017, 10 days after the positive RDT. It generally takes two days to report a culture result for *V.* cholerae [12]. Thus, the false negative culture test processes at UMTH lab led to delayed confirmation and so to delayed declaration of outbreak, which took 12 days. The late declaration, in turn, led to delayed response which might have prevented the outbreak from spreading out of Muna Garage IDP camp [13] to involve six LGAs (Fig 1B). The BMOH feared political repercussions of falsely declaring a cholera outbreak based on clinical evidence of transmission and positive RDT.

Because cholera spreads very quickly strengthening of laboratory capacity to confirm cases, in line with WHO policy [11], is urgently needed. In addition to clinical evidence of transmission and multiple positive RDTs, capacity should be increased for culture confirmation. Late setup of CTC in Muna Garage camp post declaration was a missed opportunity in appropriate response to prevent cholera spreading out of the camp.

This epidemiological setting was confounded by Eid El Kabir, a Muslim festival involving lots of food sharing and population movements that coincided with the outbreak onset, and this likely accelerated the spread of the outbreak. The threat posed by the festival might have been minimized had heightened population sensitization through community leaders and religious organizations been conducted to limit food sharing and population movements. Cholera preparedness communication’s function should include guidelines specific to handling large gatherings such as religious festivals and funerals during outbreak emergencies.

Similarly, there was a delay in evaluating sanitation issues regarding the outbreak. An investigation of latrines, in Muna garage IDP camp revealed a latrine leakage and flow to the index cases’ house; all within this household had cholera. Yet, results of this outbreak investigation influenced the response only slowly. Although authorities were alerted in a timely manner, the leaking latrine was still unrepaired one week later.

Further outbreak investigation in the IDP camps showed high-risk channels of cholera infection such as poor sanitation (overflowing non-functional and collapsed latrines, open defecation), overcrowding, and stagnant water in which children played. These high-risk channels, particularly open defecation [14] and overcrowding in the IDPs camp [15] have been linked with rapid spread of cholera [16-19]. RUWASSA was notified of these poor WASH conditions in the camp, but the response was slow. Health authorities dispatched RRTs to the field, yet their mandate and appropriateness of response was not clear. The FMWR was <u>not</u> notified of the outbreak.

In response to having not been notified of the outbreak, FMWR called on UNICEF to do more in helping the Ministries of Water Resources in carrying out their duties just as WHO does for health ministries. In this context, we think that conducting need assessments and gap analyses in Ministries of Water Resources could inform ways in which UNICEF could assist these ministries. Also strengthening intra- and inter-collaboration between water and health ministries, will improve the response. On the other hand, UNICEF-WASH’s immediate response against cholera spread was chlorination of household and sanitary facilities, which, unfortunately, led to the avoidance of all chlorinated items as rumors soon spread that chlorine was a sterilizing agent. The misconception of chlorine as a sterilizing agent was due to lack of community involvement and participation in the planning and roll-out of the chlorine response. Technical UNICEF-WASH responders entered the communities and camps without consultation with local leaders such as “Bullemas” (village headmen). Experts in communication such as UNICEF-C4D [20] were not involved either. This is an important point. Within UNICEF, there are two groups that were not communicating initially – UNICEF-WASH and UNICEF-C4D. As such the community did not know why suddenly their houses, water buckets, and latrines where being chlorinated, who the chlorinators and their intentions were, and what chlorine was. Community entry [21, 22] without community involvement and participation [23, 24] have been shown time and again in different contexts to lead to community resistance to technical interventions [25, 26], and even, to physical violence [27, 28]. When, subsequently, UNICEF-WASH involved UNICEF-C4D [20] it engaged the community and dispelled the rumors and misconception about chlorine. UNICEF-C4D involvement led to chlorine acceptance; and thus, involving communication experts in community entry is paramount to response acceptance [20].

Despite the challenges at the early phase of the outbreak BMOH, NPHCDA, and NCDC with implementing partners curtailed the outbreak with minimal fatalities. Proper and timely case management to reduce case fatality ratio (CFR = number of deaths/number of cases × 100) ensured access to ORPs or CTUs or CTCs [29-31] depending on the stage of dehydration. Overall, case management response was appropriate as ORPs where setup in hard-to-reach areas to treat less severe cases and referred severe ones to CTUs/CTCs. It achieved a 1.14% CFR, the lowest in Borno ever since outbreaks were measured [4]. Still, this is considered high as it is not below 1% as recommended by WHO [11]. A number of issues could have impacted this higher CFR including 1) late confirmation and declaration of outbreak, 2) delayed set up of CTCs in Muna Garage IDP camp (the initial thought was to manage the outbreak at normal health facilities), 3) lack of qualified people to be trained as health care workers), 4) competition between partners as to who gets the glory in treating cases, 5) patient reluctance to referrals, 6) denial of cholera by the locals, and 7) insecurity posed by Boko Haram that hindered 24 hours operation of CTUs/CTCs. The higher CFR could also have resulted from bias in the surveillance system in that the number of deaths could have been better registered than cases. Out of a total of 59 deaths, 56 died at health facility. Appropriate studies are needed to establish the links between each of these observations and increased CFR.

As an additional public health tool [11], Nigeria included a mass OCV campaign in the outbreak area for the first time ever. The vaccine was targeted at a population of 891,137 aged one year and above in six LGAs [32] to prevent the spread of the outbreak. A key success factor to the implementation of OCV was the robust polio vaccine infrastructure capacity in Borno. OCV used the mechanisms which had already been established to apply, import, and administer the vaccines rapidly, and this was accomplished in two weeks. In Iraq where an OCV campaign was carried out in response to a cholera outbreak and humanitarian crisis, the polio vaccine infrastructure was found to be a key success factor [33]. Through international networking, OCV Micro Planning used in Sierra Leon was adapted to suit the Borno context. Other success factors included the preparatory workshop in Abuja in May/June, 2017 that led to the OCV approval and subsequent registration for use in Nigeria that same year, and high advocacy with the commissioner of health publicly taking the vaccine at the onset of the campaign.

Yet, to achieve this success, the OCV team had to overcome challenges. These included difficulty in opening vials using scissors (polio vaccinators were not used to opening vials), convincing an adult population they needed the vaccines since adults thought vaccines were for children, dispelling anxiety as to reasons why the first use of OCV in Nigeria should be in Borno, limitations in cold chain capacity, obtaining data from hard-to-reach areas to inform monitoring and evaluation, and compensating vaccinators with no bank accounts. Although the polio blueprint was used, its card mode of payment was not used, but if used, may have simplified this problem. Efforts were needed to overcome controversy and rumors that came with cholera vaccine being used in Nigeria for the first time, but the life threatening cholera outbreak led people to accept the vaccine. To this end, we laud the efforts of both government, international UN partners, and NGOs for the remarkable success of the OCV campaign response.

A very critical component of the response was communications in which WHO adapted a three legged approach with focus on reaching international, national, and local communities through internet, TV, radio, banners and theater group [34]. Although, WHO communications used theater group to illustrate messages at community level, the approach lacked community input in shaping the choice of communication channels and language use, and so was rather seen as top down communication. UNICEF-C4D used a bottom up approach [20], and first mapped IDP camp and community structure to better understand the audience. They realized that Kanuri instead of Hausa was the predominant language spoken. In addition, they found that very few had access to electricity and radio let alone TV. Accordingly, the main language for messages changed from Hausa to Kanuri and the main channel for message spread became one-on-one communications. Also the role of “Bullemas” in awareness raising and community mobilization became apparent. Without downplaying the importance of top down communication, a bottom up approach was needed to understand audiences for evidence-based choices of channels and languages of communication in a resource scarce setting.

We found that the logistics team response was very appropriate in carrying out its complex operations involving many partners and flow of supplies from point of manufacture to the point of use. Whether it was setting up CTCs [31], training staff for their hygienic use, patient flow in CTCs, mapping inventory to prevent stock outs, and maintaining OCV could chain, the logistics team ensured the right products or services in the right numbers at the right place for the right price and at the right time [35]. Some challenges to the logistics team included some partners averse to releasing stock availability during inventory mapping and inability to source Ringer’s lactate locally. The limited polio infrastructure storage capacity led to trans-docking of OCV in Abuja and transport to Borno in phases, which were not cost effective. Late arrival of vaccines in Abuja caused anxiety and led to disagreements over road transport of vaccine to Borno and to chattered flights, which were costly. The chattered flight operators were put on very short notice. There were no practice exercises in cholera logistics preparedness/response plans and so loopholes in the practicality of logistics operations were unknown. This prompted the WHO-OLS unit to raise the concern that logistics was not given due attention until there was an outbreak in the state.

The coordination response lead by the BMOH was effective as we did not find gaps, overlaps, and duplications in the outbreak response between emergency responders. For instance, to ensure data harmonization and avoid double counting, partners were coordinated to record only their contributions to patients and not case counts; double counting, if allowed, would have introduced bias in CFR or the attack rate of the outbreak. However, at the <u>start</u> of the response, the coordination team had to overcome hurdles in budget allocation between partner and government, hesitancy in team formation, competing demands between insecurity and health emergency, and lack of synergy between WASH and health sectors. Key to overcoming these early phase coordination challenges was the EOC [36], which enabled the BMOH to take leadership and ownership in coordinating the emergency response. Thanks to EOC, operational data in form of epidemiological curves and geo-coordinate maps were shared between all emergency response teams at 4pm daily. We find that EOC was the hub for coordinating strategic decision making and operations related to the outbreak, which was also underscored in a systematic review [37]. Yet, there was controversy between the EOC [36, 38] and UN cluster [39] approaches to coordinating the emergency response of partners. However, this topic is beyond the scope of this study, but there is need for additional evaluation of this potential conflict.

## Conclusion/Recommendations

### Surveillance

Strengthening of culture capacity to rapidly confirm clinical and positive RDT diagnoses is urgently needed to inform outbreak declaration. While building laboratory capacity is long term and because outbreak declaration is more of a political function instead of technical, even without declaration, a full scale response should be advised given evidence of clinical transmission besides multiple positive RDTs. With timely declaration of outbreak, response would be initiated more rapidly. Some countries will not declare at all but are managing as an acute diarrheal disease. Risk communication should start immediately, and should not be delayed until declared, followed by enhanced lab/epi surveillance, case management, emergency WASH and OCV. The declaration needs to involve the local authority leaders such as “Bullemas” (village headmen).

### WASH

Clearly the WASH conditions need to improve in these IDP camps. During emergency, groups involved in WASH should partner with other agencies to ensure prompt repairs of boreholes, wells and latrines when they breakdown and emphasize the need to use latrines rather than open defecation. To avoid rumor and misconceptions, technical WASH experts should work with communication experts prior to community entry.

### Case management

The outbreak required clinical case management response to be coordinated between the groups providing clinical care and efforts were needed to avoid self-promotion among the different partners providing this care. Cholera should not be managed by a primary health care facility. Rather, timely set up of ORPs, CTUs and CTCs are highly recommended.

### OCV

The polio infrastructure was highly effective. It enabled implementation of OCV for the first time in Nigeria within two weeks. With proper training it was cost efficient, and was easily adapted to OCV peculiarities. Importantly, the card-mode of payment of staff without bank accounts used by the polio teams could also be used for OCV as well.

### Communication

An early response to an outbreak should be mapping IDP camp and surrounding community structure to inform choice of languages and channels of communication. One-on-one communication was very effective in Borno and building networks, trust, and contacts with local leaders (“Bullemas”) was paramount.

### Logistics

The regular health partners’ and logistic coordinator’s meetings was able to map supplies and maintain an inventory as part of the ongoing preparedness plans to avoid stock outs. For the future, it will be wise to improve logistics preparedness plans and make plans practical by conducting practice drills or simulations of preparedness plans to insure operational feasibility.

### Coordination

The EOC was key in strengthening coordination of the emergency response of the outbreak. Epidemiological curves and geo-coordinating map presentations during meetings at EOC at 4pm daily proved highly helpful in coordinating the response. On the other hand, coordination of WASH response was slow. To this effect, we call on RUWASSA to take a more proactive role in coordinating WASH sector response in similar emergencies in the future. This include taking ownership and leadership of WASH response in the State in close synergy with the FMWR and with UNICEF playing a critical role.

In sum, all partners should increase preparedness plans before an emergency. Partners should register (preferable through a memorandum of understanding) with the government with the understanding that they will support a coordinated response led by the government. When partners understand that the government is in charge, this facilitates government buy-in and ownership of response activities. On the other hand, government should ensure that all camps are officially recognized and establish communication channels between partners with a unified approach to accessing hard-to-reach areas. In their planning, partners need to put beneficiaries’ interest over partner interest, since the collective effort of partners is key to dealing with emergencies.

## Acknowledgements

This project could not have been possible without active support from the WHO Headquarters, and Country Office for Nigeria, Federal, State, and Local Government Area levels in Nigeria. We also acknowledge support from the DOVE-Project at Johns Hopkins University.

## Supporting information

S1 Fig. Multi-level training for the adaption of polio structure for OCV implementation.

S2 Fig. Polio card mode of payment: Could be adapted for OCV payment at ward level.

S3 Supplement. Outbreak response questionnaire.

